# CSF Boundary Disruption and Tumor-Dependent Patterns of Medulloblastoma Metastasis

**DOI:** 10.64898/2026.11.23.768674

**Authors:** Lindsay N. Ryan, Leyre Jimenez Garcia, Amy C. Gross, Resham R. Gholap, Annika M. Albrecht, Krish A. Patel, Elizabeth A. Thompson, Daphne V. Koubourli, Ryan D. Roberts, Robert J. Wechsler-Reya, James B. Reinecke

## Abstract

**Background:** Leptomeningeal metastasis is the major cause of mortality in medulloblastoma, yet the mechanisms that facilitate dissemination are poorly resolved. Here, we leveraged preclinical models to define metastatic topology, the anatomical distribution and organization of disseminated disease across craniospinal compartments and ask how topology is influenced by cerebrospinal fluid (CSF) access, primary tumor anatomy, and tumor biology.

**Methods:** Orthotopic D425 xenograft models were used to manipulate primary tumor geometry, defined by anatomical position of primary tumors within the cerebellum relative to CSF-facing boundaries. Longitudinal bioluminescence, survival, primary growth architecture at the tumor– host interface, CSF-boundary breach, and craniospinal metastatic distribution were quantified. Additional medulloblastoma models MED411, MED2112, and CHLA-01-MED were implanted at a common midline cerebellar location to assess tumor-intrinsic contributions to metastatic topology.

**Results:** While primary tumor geometry altered disease progression and survival, it did not significantly alter metastasis distribution. Strikingly, growth architecture and primary tumor burden correlated with dissemination. Despite their large size, circumscribed tumors frequently remained localized, whereas smaller, disruptive primary growth architecture was strongly associated with boundary breach, intracranial, and spinal dissemination. Conversely, when anatomical origin was held constant, distinct medulloblastoma models generated significantly different metastatic topologies.

**Conclusions:** Disruption of CSF-facing anatomical boundaries is associated with distant dissemination, whereas the distribution of established metastatic disease differs substantially among medulloblastoma models despite standardized implantation conditions. These findings distinguish anatomical factors associated with dissemination from tumor-dependent patterns of metastatic colonization and establish an experimental framework for investigating the mechanisms underlying metastatic topology.

**Key points:**

- Leptomeningeal dissemination occurs across primary tumor geometries, but extent of metastatic disease is heterogeneous
- Primary tumor architecture and growth is closely tied to leptomeningeal dissemination, with CSF-boundary breach as an important determinant
- Tumor-intrinsic biology influences preferential patterns of metastatic dissemination

**Importance of the Study:** Medulloblastoma metastasis is typically characterized by the presence and extent of disseminated disease, with limited consideration of its precise anatomical distribution with respect to diverse cellular niches throughout the craniospinal axis. Here, we demonstrate that dissemination is strongly associated with disruption of CSF-facing anatomical boundaries, whereas the distribution of established metastases is strongly dependent on tumor model identity. These findings establish metastatic topology as a distinct feature of medulloblastoma dissemination and suggest that anatomical access and site-specific colonization are governed by separable biological processes.

**Lay Summary:** Medulloblastoma is an aggressive childhood brain tumor. Although modern treatments cure many children, the disease returns to up to 30% of patients. Medulloblastoma can also spread beyond its original location through the fluid surrounding the brain and spinal cord, a process called leptomeningeal metastasis. This spread is associated with poor outcomes, yet scientists do not fully understand why some tumors spread or what determines where metastatic tumors develop.

In this study, we used mouse models of medulloblastoma to investigate how the location and growth characteristics of the original tumor influence metastatic spread. We found that metastasis was strongly associated with tumor disruption of the natural barriers separating brain tissue from fluid-filled spaces, rather than simply the amount of tumor growth. Importantly, different medulloblastoma models developed distinct patterns of metastasis throughout the brain and spinal cord, even when the original tumors were implanted in the same location.

These findings suggest that the ability of tumors to access fluid pathways and their tendency to colonize particular locations are influenced by different factors. Understanding these processes may help researchers develop new strategies to prevent or treat metastatic medulloblastoma.

## Introduction

Medulloblastoma is the most common malignant brain tumor of childhood and comprises four molecular groups with distinct developmental origins, molecular drivers, and clinicopathological behaviors[1–8]. Although advances in surgery, craniospinal irradiation, and multi-agent chemotherapy have substantially improved survival, metastatic progression remains a dominant cause of mortality[9–15]. Despite the central role of metastasis in determining outcome, the factors that govern how medulloblastoma cells access and colonize distinct compartments of the craniospinal axis remain poorly understood.

Clinically, disseminated medulloblastoma is staged using the Chang criteria: M0 indicates no evidence of metastatic disease, M1 positive cerebrospinal fluid (CSF) cytology without radiographically detectable metastasis, M2 intracranial metastatic disease, M3 spinal metastatic disease, and M4 extracranial spread[16]. Although this framework effectively captures the broad extent of dissemination, it does not reflect the marked spatial heterogeneity that exists within individual metastatic stages. M2 disease may include intraventricular metastases, bulky disease within intracranial cisterns, or diffuse leptomeningeal spread over supratentorial or cerebellar surfaces. Similarly, M3 disease can manifest as thin, laminar surface coating or nodular leptomeningeal deposits along the spinal cord, nerve-root-associated disease, or bulky tumor within the distal thecal sac. We refer to this spatial organization of disseminated disease across the craniospinal axis as metastatic topology. Defining the determinants of metastatic topology may reveal biologically distinct modes of dissemination that are obscured by broad metastatic stage alone.

Primary medulloblastoma location is heterogeneous and non-randomly associated with molecular groups. SHH-activated tumors frequently arise within the cerebellar hemispheres, whereas Group 3 and Group 4 tumors more commonly arise in midline/posterior fossa locations closely associated with the fourth ventricle[17]. These groups also differ substantially in metastatic propensity and patterns of relapse. Group 3 and Group 4 tumors are frequently metastatic at diagnosis and commonly relapse with distant disease, whereas SHH-activated tumors are less frequently metastatic at presentation and more often recur locally[14, 18, 19]. Thus, primary tumor anatomy and tumor-intrinsic biology co-vary clinically, making their relative contributions to metastatic dissemination difficult to disentangle.

The anatomy of the posterior fossa provides several distinct interfaces through which a growing tumor may encounter the CSF compartment[20]. Superficial cerebellar tumors directly approach the pial and subarachnoid surfaces, whereas deeply situated lateral tumors lie closer to the lateral recesses of the fourth ventricle and adjacent posterior fossa cisterns. Midline tumors are positioned adjacent to the fourth ventricle, which is continuous caudally with the cisterna magna and spinal subarachnoid space and rostrally with the cerebral ventricular system through the aqueduct. These anatomical relationships could constrain the initial routes available to disseminating tumor cells [21].

In this study, we used stereotactic implantation to establish medulloblastoma tumors at distinct locations within the mouse cerebellum and thereby experimentally vary primary tumor geometry relative to CSF-facing boundaries. We quantified disease progression, survival, primary tumor growth architecture, CSF-boundary breach, and craniospinal metastatic distribution. When tumor identity was held constant, primary geometry altered disease course but did not significantly impact metastasis. However, primary tumor architecture was strongly associated with boundary breach and dissemination. Conversely, when primary geometry was held constant, distinct medulloblastoma models generated markedly different patterns of craniospinal colonization despite comparable anatomical opportunity. Collectively, these findings suggest that metastatic topology is a multifactorial process influenced by both anatomical opportunity and tumor biology.

## Materials and Methods

### Cell lines

Cell lines utilized in this manuscript were routinely authenticated with short tandem repeat profiling and *Mycoplasma* testing (Laboratory Corporation of America). Medulloblastoma cell lines used in this study include: D425 (Group 3), MED-411FH (Group 3), MED-2112FH (Group 3), and CHLA-01-MED (Group 4). D425 Med (D425) cells (RRID: CVCL_1275) were derived from a primary medulloblastoma tumor resection at Duke University Medical Center and obtained from the laboratory of Dr. Jeffery Leonard[22]. Med-411FH (MED411) cells and Med-2112FH (MED2112) cell lines derived from early passage patient-derived xenografts developed by the laboratory of Dr. Jim Olson and distributed by the Brain Tumor Resource Lab at Seattle Children’s Hospital[23]. CHLA-01-MED was obtained from American Type Culture Collection[24]. Cell lines were maintained as previously described[25]

### Animal studies

All animal procedures were approved by the Institutional Animal Care and Use Committee of the Abigail Wexner Research Institute at Nationwide Children’s Hospital (IACUC protocol AR15-00022). CB17-SCID mice [CB17SC scid □−□/□−□; RRID: IMSR_TAC:cb17sc] were used for all *in vivo* experiments.

#### Evans blue validation

6–8-week-old mice were anesthetized with isoflurane and positioned in a stereotaxic apparatus (Kopf Instruments, Model 940). Mice received a 0.5 µL intracerebellar injection of Evans Blue (0.5 mg/mL in PBS; Sigma-Aldrich, E2129-10G) at one of three anatomically defined orthotopic locations: deep (2 mm lateral, 2 mm posterior, and 2.5 mm ventral relative to lambda), midline (0.5 mm lateral, 2 mm posterior, and 2.5 mm ventral relative to lambda), or shallow (2 mm lateral, 2 mm posterior, and 1.75 mm ventral relative to lambda). Fifteen minutes following Evans blue injection, mice were euthanized, a sternotomy was performed, and transcardial perfusion was conducted via the left ventricle using 20 mL of 0.9% saline (BD PosiFlush; Cat. No. 306546), followed by 30 mL of neutral buffered formalin (NBF; Sigma-Aldrich, Cat. No. HT5012-1CS). Skulls were dissected, cleared of connective tissue and placed in NBF at 4 °C for 48 h. Brains were dissected from the skull and cut into 1 mm coronal slices after NBF incubation.

#### Spontaneous Leptomeningeal Metastasis Model

6–8-week-old mice were anesthetized with isoflurane and placed in a stereotaxic apparatus (Kopf Instruments, Model 940). Intracerebellar injections of 8 × 10^4 medulloblastoma cells in 3 µL of plain DMEM media were performed at three anatomically defined orthotopic locations as determined by Evans blue validation: deep (2 mm lateral, 2 mm posterior, and 2.5 mm deep relative to lambda), midline (0.5 mm lateral, 2 mm posterior, and 2.5 mm deep relative to lambda), and shallow (2 mm lateral, 2 mm posterior, and 1.75 mm deep relative to lambda). Post-operative care included analgesia and daily monitoring. Mice were euthanized at neurological or humane endpoints, defined by rapid weight loss, an enhanced body condition score of <9, and/or neurological signs including head tilt, seizures, or log rolling. Following euthanasia, tissues were collected and processed for histology

### Histology

Upon reaching humane endpoint criteria, mice were euthanized, a sternotomy was performed, and mice were perfused as previously described[25]. Heads and spinal columns were dissected, cleared and dropped in neutral-buffered formalin at 4 °C for 48 h. After fixing, brains were dissected from the skull, and spines were decalcified in 10% EDTA (Elabscience, E-IR-R112) for 10 days at room temperature and then processed for paraffin embedding (FFPE). FFPE sections were counterstained with hematoxylin and eosin (H&E) via conventional methods for histological analysis.

### Thick Tissue Clearing, Confocal and light microscopy

Brain slices of 1 mm thickness were stained with Wheat Germ Agglutin conjugated to Alex Fluor 488 and DAPI overnight and cleared using Ce3D method (BioLegend cat #427703, [26]). Confocal microscopy images were obtained using a Crest-X-Light V3 spinning disk confocal on a Nikon Ti2-E inverted microscope, equipped with a Hamamatsu ORCA Fusion camera. Raw ND2 files were imported into ImageJ for post-image processing and quantitation. All manipulation of images (brightness/contrast) was done uniformly to images. FFPE sections stained with H&E were imaged with the Aperio FL ScanScope digital slide scanner at 40x magnification.

### Image analysis: Tumor geometry, architecture and dissemination

Primary tumor growth architecture was classified on hematoxylin and eosin (H&E)-stained sections into three categories: circumscribed (0), focally disruptive (1), and disruptive (2), based on the morphology of the tumor–host interface. Tumor solidity was quantified from segmented primary tumor boundaries in ImageJ as the ratio of tumor area to the area of its convex hull. Primary tumor burden was assessed using a semiquantitative ordinal scale ranging from 0 to 3 (absent, limited-25% of posterior fossa, moderate-up to 50% of posterior fossa, and extensive->50% involvement and/or significant tissue disruption/mass effect). Given the limitations of FFPE thin sections, histological primary burden scores were not considered equivalent to volumetric tumor measurements. Disruption of CSF-facing anatomical boundaries was assessed separately, including the cerebellar pia, transverse fissure, fourth ventricular interface, and lateral recesses (Table 1, glossary of definitions).

**Table 1:** Glossary for terminology used when describing defining characteristics of primary tumors.

| **Term** | **Definition** | **Scoring criteria** |
|---|---|---|
| Tumor geometry | The anatomical placement of a primary tumor within the cerebellum | Categorical, based on predetermined cerebellar locations on the x, y and z plane. |
| Growth architecture | The pattern by which a primary tumor grows within the cerebellum in relation to the surrounding non-tumor stroma | Ordinal, number scoring from 0 to 2 based on histological analysis of primary tumor in 2D. |
| CSF-boundary breach | The disruption by a primary tumor of a CSF interface separating the primary tumor from a CSF-containing space and disrupting the parenchymal and/or pial boundary. | Binary, number scoring of 0 or 1 based on the absence or presence of CSF interface disruption in histology. |
| Solidity | Quantitative measure of the compactness of a primary tumor based on the ratio of the tumor volume to the volume of its convex hull | Quantitative, based on total primary tumor area over the area of the smallest convex shape that enclosed the tumor. |

Metastatic disease was evaluated across nine anatomical topology features: cranial surface leptomeningeal involvement, basal/ventral cisternal disease, fourth ventricular occupancy or local extension, rostral ventricular disease (third and lateral ventricles), spinal surface involvement, rostrocaudal spinal surface extent, nerve-root/cauda burden, intrathecal burden, and dorsal-root-ganglion involvement. Each feature was assigned an ordinal score from 0 to 3 (absent, limited-to individual cells, moderate-multi-focal disease, or extensive-diffuse laminar or space occupying lesions). The anatomical distribution and burden of metastatic disease was assessed by serial sectioning of brains in coronal or sagittal orientation. Spines were first cut into cervical, thoracic, lumbar components, hemisected and then serial sectioned in the sagittal plane.

Distant intracranial dissemination was defined by evidence of cranial surface, basal/ventral cisternal, or rostral ventricular disease. Fourth ventricular occupancy alone was excluded from this binary endpoint because it could represent contiguous extension from the primary tumor. Spinal dissemination was defined by involvement of the spinal surface, nerve roots/cauda equina, intrathecal compartment, or dorsal root ganglia.

Composite intracranial extent was calculated as the sum of cranial surface, ventral cisternal, and rostral ventricular scores. Composite spinal surface extent was calculated as the sum of spinal surface burden and rostro-caudal spinal surface extent (possible range, 0–6). Nerve-root, intrathecal, and DRG involvement were retained as distinct topology features rather than incorporated into the spinal surface composite. A score of zero represented an evaluated compartment without detectable disease, whereas missing values indicated tissue or compartments that could not be adequately evaluated.

### Statistical analysis

Statistical analyses and data visualization were performed using R (4.5.3) or Graph Pad Prism (v10). R packages included readxl, dplyr, ggplot2, cluster, and vegan. Individual animals were treated as independent biological replicates.

Differences in the incidence of intracranial and spinal dissemination were evaluated using Fisher’s exact tests. Continuous and ordinal measurements were compared across groups using Kruskal–Wallis tests with post-hoc analysis using Wilcoxon rank tests with Bonferroni correct for multiple comparisons where applicable. Associations between primary tumor burden and metastatic extent were assessed using Spearman rank correlation coefficients. Analyses were performed using all evaluable animals and, where indicated, restricted to animals with established dissemination.

An exploratory geometry-stratified permutation analysis evaluated the relationship between primary tumor burden and dissemination while preserving the observed implantation-geometry groups. Dissemination labels were randomly reassigned within each geometry group, and a within-group-centered association statistic was recalculated over 9,999 permutations to derive two-sided permutation p-values.

Global differences in metastatic topology were evaluated using permutational multivariate analysis of variance (PERMANOVA). Pairwise Gower dissimilarities were calculated from the selected semiquantitative anatomical scores using the daisy function in the cluster package. Ordinal scores were entered as numeric values and range-standardized in the Gower calculation. PERMANOVA was performed using adonis2 in vegan with 9,999 permutations. Homogeneity of multivariate dispersion was evaluated using betadisper and permutest. Principal coordinates analysis was used for exploratory visualization of the resulting dissimilarity matrix. All reported tests were two-sided, with p < 0.05 considered statistically significant. Descriptive results are presented as individual observations, frequencies, medians, and interquartile ranges, or other summaries as appropriate for the measurement scale. Sample sizes and exclusions are reported for individual analyses.

## Results

### Stereotactic modeling establishes workflow for generating distinct primary tumor–CSF geometries in the mouse posterior fossa

Medulloblastoma dissemination is clinically categorized using M-stage criteria that distinguish cytologic, intracranial, spinal, and extra-neural disease[16, 27]. Although this framework provides an important measure of metastatic extent, these broad M-stage categories encompass substantial spatial heterogeneity. Intracranial metastatic disease may involve the cerebral or cerebellar surface, basal cisterns, or ventricular system, whereas spinal disease may present as diffuse surface coating, nerve-root/cauda involvement, or bulky intrathecal lesions. We therefore use the term metastatic topology to describe the anatomical distribution of disseminated tumor across these craniospinal compartments (Figure 1A).

**Figure 1:**
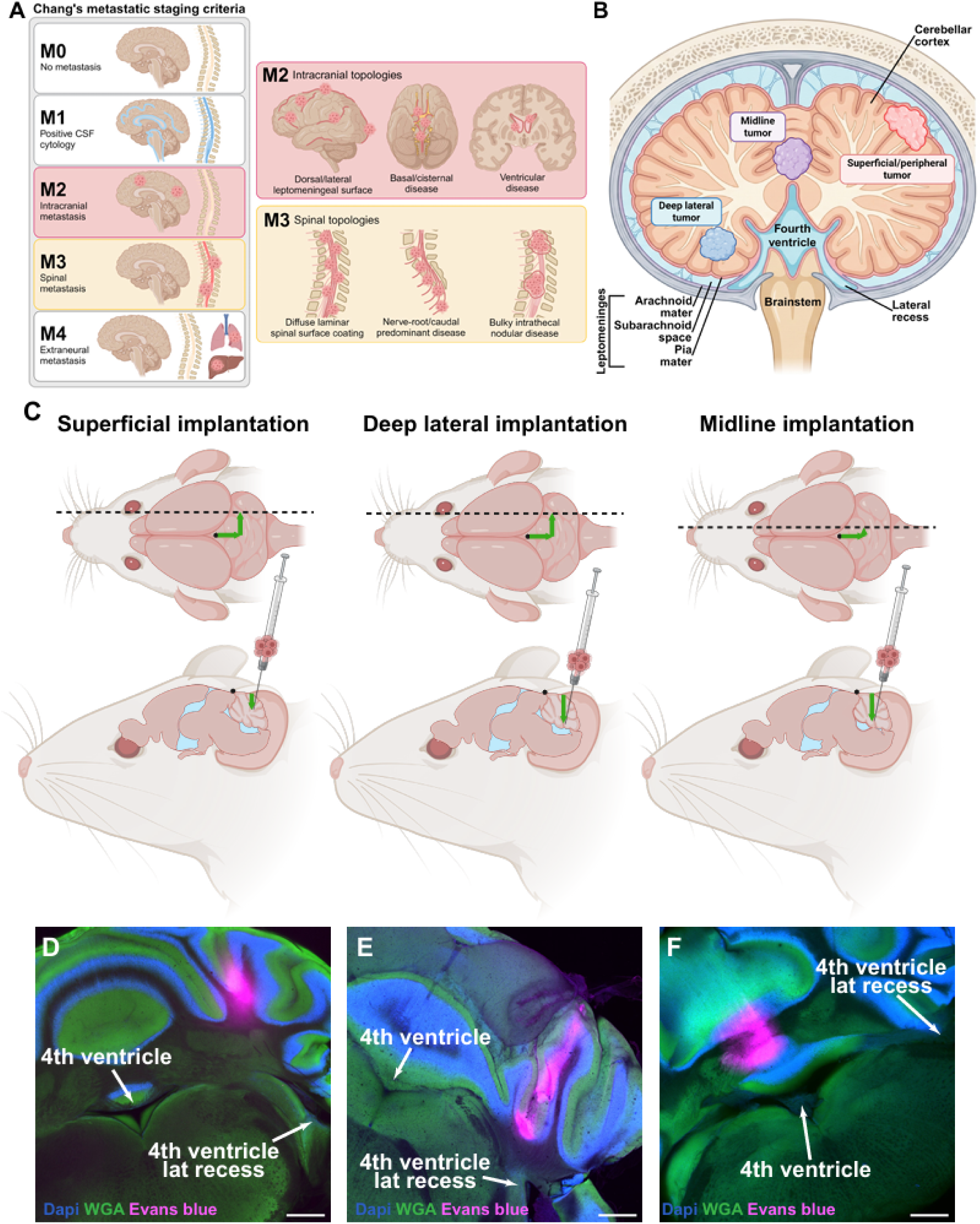
Modeling primary tumor geometry heterogeneity in mice to study metastasis topology. A) schematic representation of the Chang criteria used for staging dissemination in patients. Created in BioRender. Jimenez Garcia, L. (2026). B) Schematic of cerebellar anatomy illustrating position of medulloblastoma primary tumors relative to CSF boundaries. Created in BioRender. Jimenez Garcia, L. (2026). C) Experimental outline for stereotactic injection into distinct anatomical regions of mouse cerebellum. Created in BioRender. Jimenez Garcia, L. (2026). D-F) Representative images of cleared mouse cerebellum following injection of Evans Blue (red) at defined coordinates (see methods). Wheat germ agglutin (WGA; green) and DAPI (blue) highlight cerebellar anatomy. Scale bar 100 µm.

Because the cerebellum contains multiple anatomically distinct interfaces with the cerebrospinal fluid compartment, we asked whether the geometry of the primary tumor relative to these interfaces could influence subsequent dissemination. Tumors arising near the cerebellar surface are positioned adjacent to the pial/subarachnoid space, whereas deeper lateral tumors may encounter the lateral recess of the fourth ventricle or superficial pial boundaries during growth. Midline tumors occupy a distinct relationship relative to the fourth ventricle (Figure 1B). We hypothesized that these differences in primary tumor geometry would create distinct opportunities for CSF access and thereby influence metastatic behavior.

To experimentally separate primary tumor geometry from tumor-cell identity, we established stereotactic coordinates that reproducibly positioned the same medulloblastoma model at superficial, deep-lateral, or midline sites within the mouse cerebellum (Figure 1C). Small volume superficial injections localized within the cerebellar hemisphere immediately adjacent to the pial surface, deep-lateral injections were positioned near the lateral recess of the fourth ventricle, and midline injections localized within the cerebellar vermis adjacent to the fourth ventricular compartment. Fluorescent tracer studies confirmed localization of the initial injection deposits adjacent to the intended CSF-facing boundaries without gross disruption of these interfaces (Figure 1D-F). We next used these defined geometries to determine whether primary anatomical position influenced tumor progression and metastatic dissemination.

### Primary tumor geometry is associated with differences in time to neurological endpoint without detectable differences in bioluminescent tumor burden

Luciferase-expressing D425 medulloblastoma cells were implanted into the cerebella of immunodeficient mice to evaluate the influence of primary tumor geometry on disease progression. Tumor growth was monitored longitudinally by bioluminescence imaging (BLI), and animals were euthanized upon reaching clinical or neurological endpoints. Brain and spinal tissues were subsequently collected for histopathological assessment of primary tumor growth and disseminated disease (Figure 2A).

**Figure 2:**
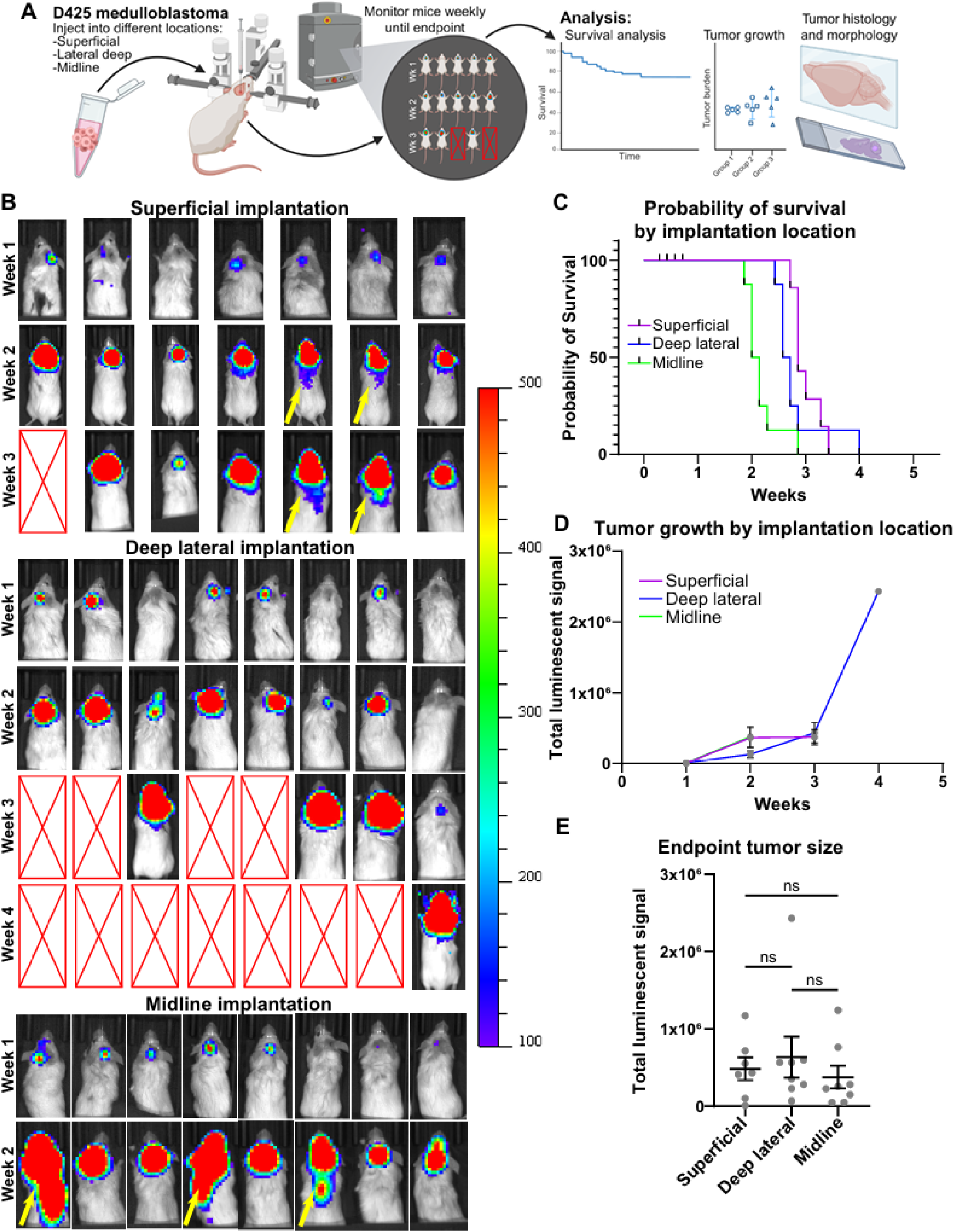
Effect of primary tumor geometry on tumor progression and survival. A) Schematic of experimental approach. Created in BioRender. Jimenez Garcia, L. (2026). B) Representative images of tumor progression by bioluminescence. Arrows denote signal consistent with spinal disease. C) Kaplan-Meier survival plot by implantation location. D-E) Quantitation of bioluminescence over time and at neurological endpoint. Error bars represent mean and standard error of the mean. Ordinary one-way ANOVA with Tukey’s multiple comparisons test, ns = not significant.

Progressive intracranial bioluminescent signal was observed across all three implantation cohorts, with detectable spinal signal in 2/7 superficial, 0/8 deep lateral, and 3/8 midline animals (Figure 2B, arrows). Mice in the midline cohort reached neurological endpoint at 15.1 ± 2.2 days, compared with 20.9 ± 2.0 and 19.6 ± 3.5 days in the superficial and deep lateral cohorts, respectively (Figure 2C). Despite these differences in time to endpoint, longitudinal bioluminescent signal and terminal BLI measurements did not differ significantly between implantation groups (Figures 2D–E, Ordinary one-way ANOVA with Tukey’s multiple comparisons test, ns = not significant).

These findings suggest that primary tumor geometry may influence the timing of disease progression to neurological endpoint without corresponding differences in BLI-detectable tumor burden. However, because bioluminescence may incompletely capture diffuse leptomeningeal disease, comparable imaging signals cannot establish equivalent total disease burden. We therefore performed a comprehensive histopathological assessment of primary tumor architecture, CSF-boundary involvement, and craniospinal dissemination to better characterize the anatomical determinants of disease progression.

### Leptomeningeal dissemination occurs across primary tumor geometries with heterogeneous metastatic extent

To determine whether primary tumor geometry influences the anatomical distribution and extent of metastatic disease, we performed comprehensive histopathological assessment of cranial and spinal tissues from D425-bearing mice across the three implantation cohorts. Representative sections demonstrated substantial heterogeneity in metastatic involvement, ranging from limited leptomeningeal deposits to extensive tumor coating of the craniospinal surfaces (Figure 3A). Although extensive spinal involvement was observed more frequently among individual midline and deep lateral tumors, leptomeningeal dissemination occurred across all three implantation geometries.

**Figure 3:**
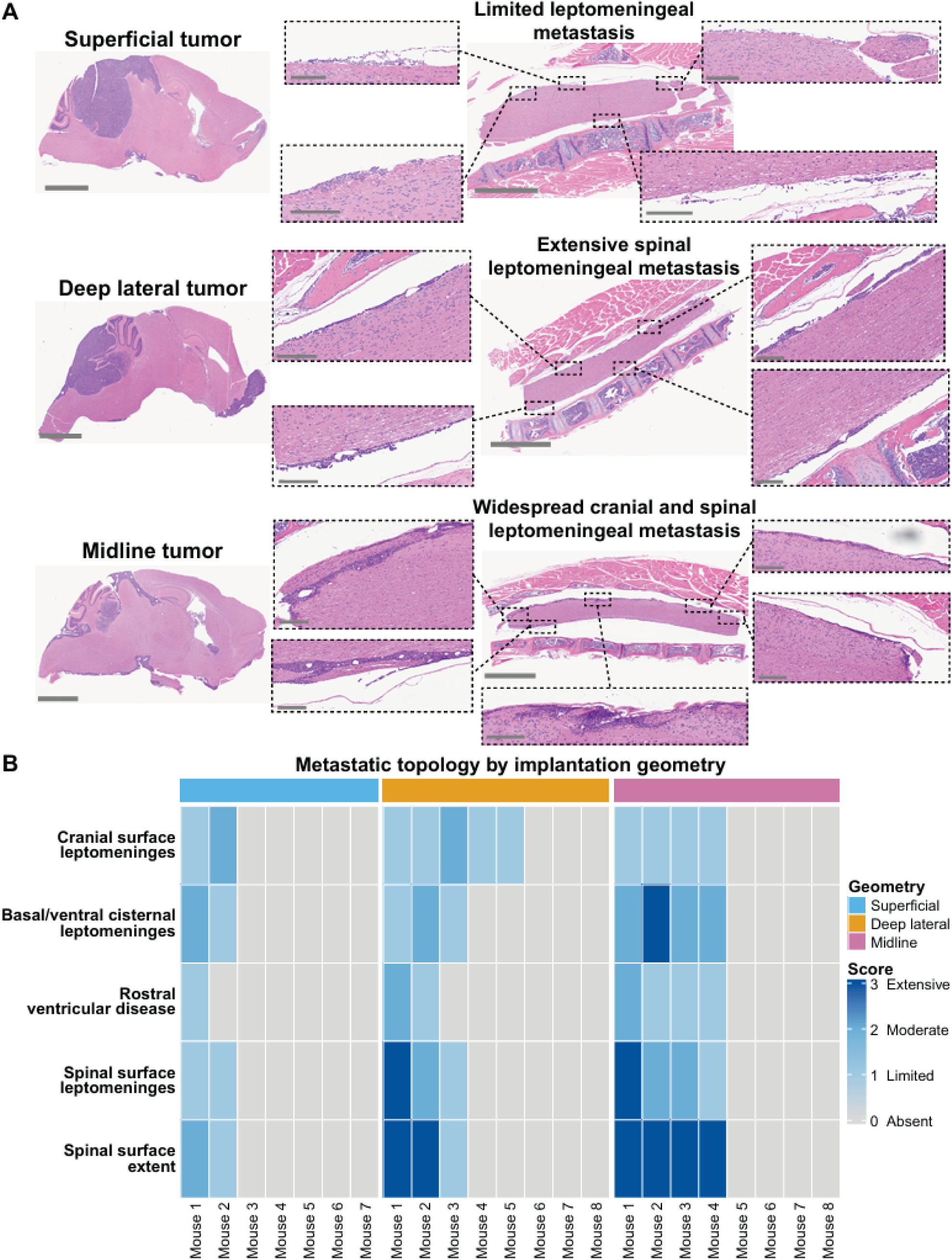
Role of primary tumor geometry on metastatic topology. A) Representative histological images of primary tumor and spinal metastasis. Scale bar = 2 mm. Inset scale bar = 200 µm. B) Heatmap portraying ordinal scoring for metastatic topology by individual mouse per geometry cohort.

The incidence of intracranial dissemination did not differ significantly between superficial (2/7), deep lateral (5/8), and midline (4/8) cohorts (Supplemental Figure 1A, Fisher’s exact test, p = 0.508). Similarly, spinal dissemination was observed in 2/7 superficial, 3/8 deep lateral, and 4/8 midline animals (Supplemental Figure 1B, Fisher’s exact test, p = 0.866). To further characterize differences in disease distribution, we scored metastatic involvement across predefined craniospinal compartments using an ordinal histopathological scale (Figure 3B). Despite apparent differences in the extent of spinal surface involvement among individual animals, neither intracranial nor spinal metastatic extent differed significantly across implantation geometries. Multivariate analysis likewise failed to identify significant separation in overall metastatic topology.

Together, these findings indicate that primary tumor geometry alone does not reliably predict the occurrence or anatomical distribution of leptomeningeal dissemination, suggesting that additional determinants govern metastatic progression.

### Primary tumor growth architecture and CSF-boundary breach are associated with leptomeningeal dissemination

Previous MRI-based studies have demonstrated subgroup-associated differences in medulloblastoma primary tumor localization and growth characteristics, including tumor size, shape, and margin definition[2, 6, 17, 19]. Although metastatic disease can occur in patients with relatively small primary tumors, the relationship between primary tumor architecture, local tissue invasion, and leptomeningeal dissemination remains incompletely understood. Moreover, radiographic assessment provides limited resolution of the tumor–host interface and microscopic involvement of CSF-facing anatomical boundaries. We therefore reasoned that detailed histopathological characterization of primary tumor growth architecture, paired with compartment-specific assessment of microscopic metastatic disease, would provide greater insight into the anatomical features associated with dissemination.

We identified three distinct primary tumor growth architectures: circumscribed, focally disruptive, and disruptive (Figure 4A). Circumscribed tumors formed cohesive masses with smooth, well-defined tumor–host interfaces. Focally disruptive tumors exhibited predominantly circumscribed growth with localized irregularity at the tumor margin, whereas disruptive tumors demonstrated more extensively irregular growth patterns. These morphological distinctions were supported by quantitative assessment of tumor solidity (Figure 4B-C). Interestingly, disruptive tumors frequently exhibited lower histologically assessed primary tumor burden than either circumscribed or focally disruptive tumors (Figure 4D, median ordinal burden scores of 1, 3, and 3, respectively; Kruskal–Wallis p = 0.014). Five of eight disruptive tumors exhibited the lowest primary burden score, compared with none of the 15 circumscribed or focally disruptive tumors (Figure 4E). These observations suggest that extensive disruption of the tumor–host interface does not necessarily require the development of a large, cohesive primary tumor mass. Although primary tumor architecture varied across implantation geometries, the association did not reach statistical significance (Supplementary Figure 2A).

**Figure 4:**
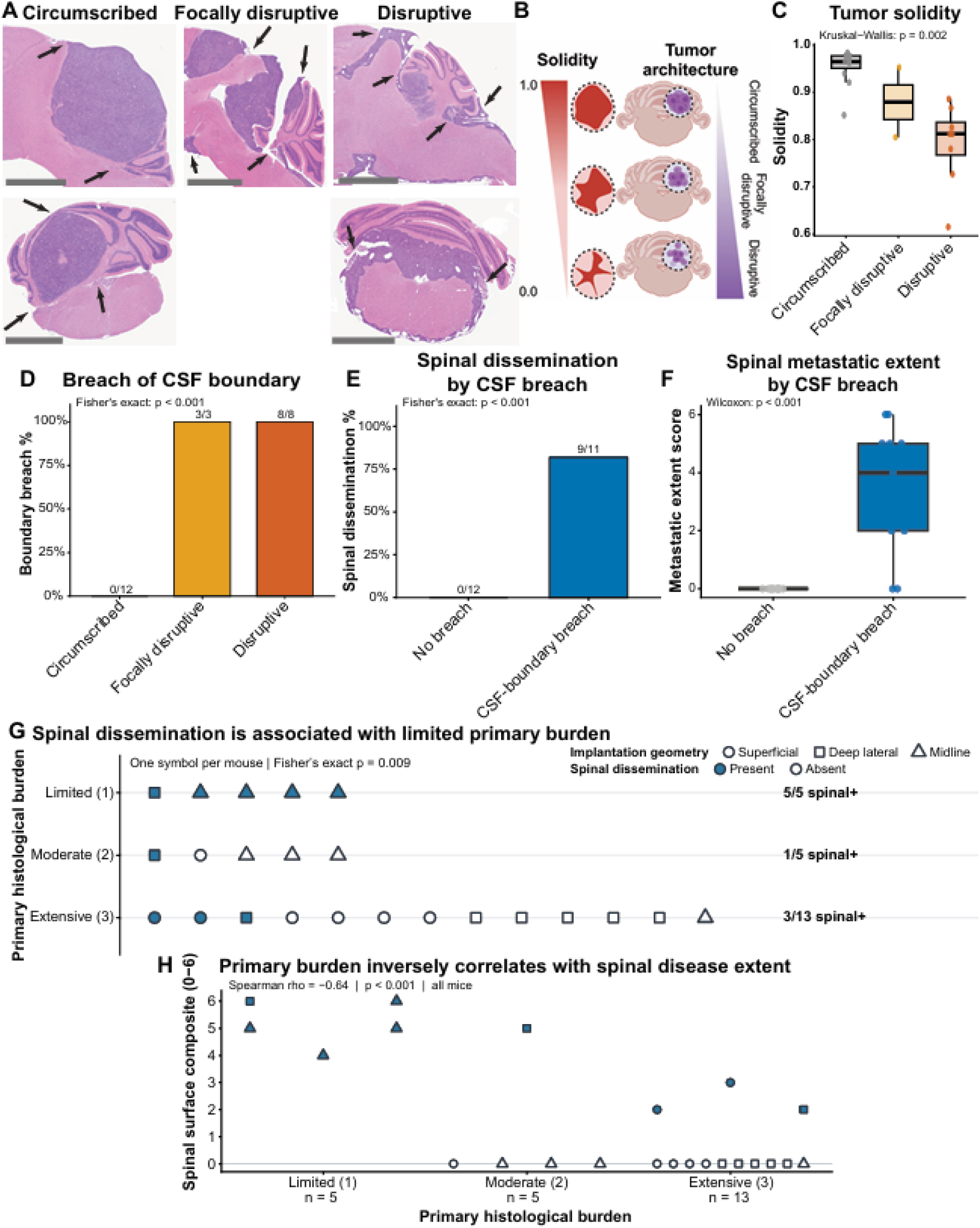
Relationship between primary tumor architecture, burden, and dissemination. A) Representative histological images of tumor architectures. Scale bar = 2 mm. Inset scale bar = 200 µm. B-C) Schematic and quantitation of tumor solidity by architecture. Created in BioRender. Jimenez Garcia, L. (2026). D-F) Relationship between architecture, CNS barrier breach, and dissemination. G-H) Spinal dissemination and extent of spinal dissemination by primary tumor burden.

Histopathological examination further revealed distinct patterns of local tissue invasion and CSF-boundary involvement. Superficial and deep lateral tumors demonstrated the capacity to disrupt cerebellar pial surfaces and extend into the adjacent longitudinal fissure between the cerebellum and cerebrum, as well as involve the fourth ventricular and lateral recess compartments. In contrast, boundary disruption among midline tumors was restricted to the fourth ventricular interface. Given the two-dimensional nature of histological sections, which precluded reliable estimation of total primary tumor volume, subsequent analyses focused on growth architecture, semiquantitative primary burden, anatomical boundary involvement, and metastatic distribution.

We next examined whether primary tumor architecture and histological disruption of CSF-facing boundaries were associated with metastatic dissemination. Circumscribed tumors maintained intact anatomical boundaries and demonstrated no evidence of leptomeningeal dissemination. In contrast, tumors exhibiting CSF-boundary breach frequently developed craniospinal metastatic disease. Notably, spinal dissemination was absent in all 12 animals without histological evidence of boundary breach but was identified in 9 of 11 animals with boundary disruption (p < 0.001; Figure 4F-H). Together with the observation that disruptive tumors frequently exhibited limited primary tumor burden, these findings suggest that metastatic dissemination is more closely associated with disruption of CSF-facing anatomical boundaries than with the accumulation of a large primary tumor mass.

We further explored whether the anatomical site of boundary disruption was associated with the extent of metastatic disease. Among tumors with CSF-boundary breach, involvement restricted to cerebellar pial or fissural surfaces was associated with less extensive spinal disease than breach involving the fourth ventricular or lateral recess compartments (Supplemental Figure 2B-C). Although spinal dissemination occurred in 1 of 3 animals with pial/fissural-only breach compared with 8 of 8 animals with ventricular/lateral-recess involvement, the difference in incidence did not reach statistical significance (p = 0.055). Spinal metastatic extent nevertheless differed between these groups (p = 0.022), suggesting that the anatomical route of CSF access may contribute to the extent of subsequent dissemination.

Collectively, these findings identify primary tumor growth architecture and CSF-boundary disruption as important anatomical correlates of leptomeningeal dissemination. Importantly, extensive boundary disruption and metastatic spread were observed even among tumors with limited primary burden, suggesting that dissemination is not simply a consequence of increasing primary tumor mass. Having identified primary tumor characteristics associated with dissemination, we next investigated whether distinct medulloblastoma models exhibit characteristic metastatic topologies despite shared anatomical starting conditions.

### Tumor identity is associated with distinct metastatic topologies despite shared implantation geometry

Recent developmental and molecular studies have demonstrated that Group 3 and Group 4 medulloblastomas share overlapping transcriptional programs and developmental origins within the human rhombic lip[1, 2, 6]. Consistent with these findings, clinical imaging studies have demonstrated that tumors from both subgroups predominantly localize to the cerebellar midline, frequently involving or expanding into the fourth ventricle[2, 17]. Despite these shared anatomical characteristics, Group 3 and Group 4 medulloblastomas exhibit considerable heterogeneity in the distribution and morphology of leptomeningeal metastases[18, 19, 28]. We therefore reasoned that comparing distinct medulloblastoma models arising from a common anatomical starting location would allow us to investigate the contribution of tumor identity to metastatic topology.

To test this hypothesis, we generated midline orthotopic tumors using three additional patient-derived medulloblastoma models: MED411 and MED2112, both derived from MYC-amplified Group 3 tumors, and CHLA-01-MED, a Group 4 model. Animals were monitored longitudinally and euthanized upon reaching neurological endpoint. Time to endpoint varied substantially between models, with MED411 and MED2112 tumors reaching endpoint within 28–31 and 26– 29 days, respectively, whereas CHLA-01-MED demonstrated considerably prolonged survival, reaching endpoint between 101 and 123 days following implantation (Supplementary Figure 3A).

Histopathological examination revealed substantial differences in primary tumor growth characteristics despite shared midline implantation geometry. MED2112 tumors consistently formed small, irregularly shaped primary masses with extensive disruption of the fourth ventricular interface. Interestingly, when MED2112 cells were implanted using shallow lateral cerebellar coordinates, all three animals developed large, circumscribed primary tumors without histological evidence of CSF-boundary breach, in contrast to the smaller, disruptive tumors observed in all five midline animals (Supplementary Figure 3B). Although this comparison was not powered to establish statistically significant differences in growth or dissemination, it suggested that anatomical context could influence primary tumor growth architecture even within an individual medulloblastoma model.

MED411 tumors similarly exhibited relatively limited parenchymal primary tumor burden, although their growth architectures were more heterogeneous, encompassing including both focally disruptive and disruptive morphologies (Supplemental Figure 3C). In contrast, CHLA-01-MED tumors developed large, expansile primary masses that filled and distorted the fourth ventricle without a clearly defined cerebellar parenchymal component (Supplemental Figure 3D). Thus, despite a shared anatomical starting position, the models exhibited distinct patterns of primary tumor growth and ventricular involvement. Having established the primary tumor characteristics of these additional models, we next examined whether tumor identity was associated with distinct patterns of metastatic colonization across the craniospinal axis.

Leptomeningeal metastases can occupy anatomically distinct compartments throughout the craniospinal axis, including the ventricular system, cranial and spinal leptomeninges, nerve roots, and intrathecal spaces. Whether the distribution of metastatic disease is primarily determined by the anatomical location of CSF access or reflects characteristics of individual tumors remains unclear. To address this question, we compared the anatomical distribution of metastatic disease across four medulloblastoma models established using a common midline implantation geometry.

Despite shared implantation coordinates, the models exhibited markedly different patterns of craniospinal disease (Figure 5A–D). D425 tumors frequently demonstrated diffuse spinal surface involvement, whereas MED411 and MED2112 exhibited more prominent involvement of additional spinal compartments, including nerve roots and intrathecal spaces. CHLA-01-MED demonstrated a distinct pattern characterized by substantial ventricular involvement, including local extension into the fourth ventricular compartment.

**Figure 5:**
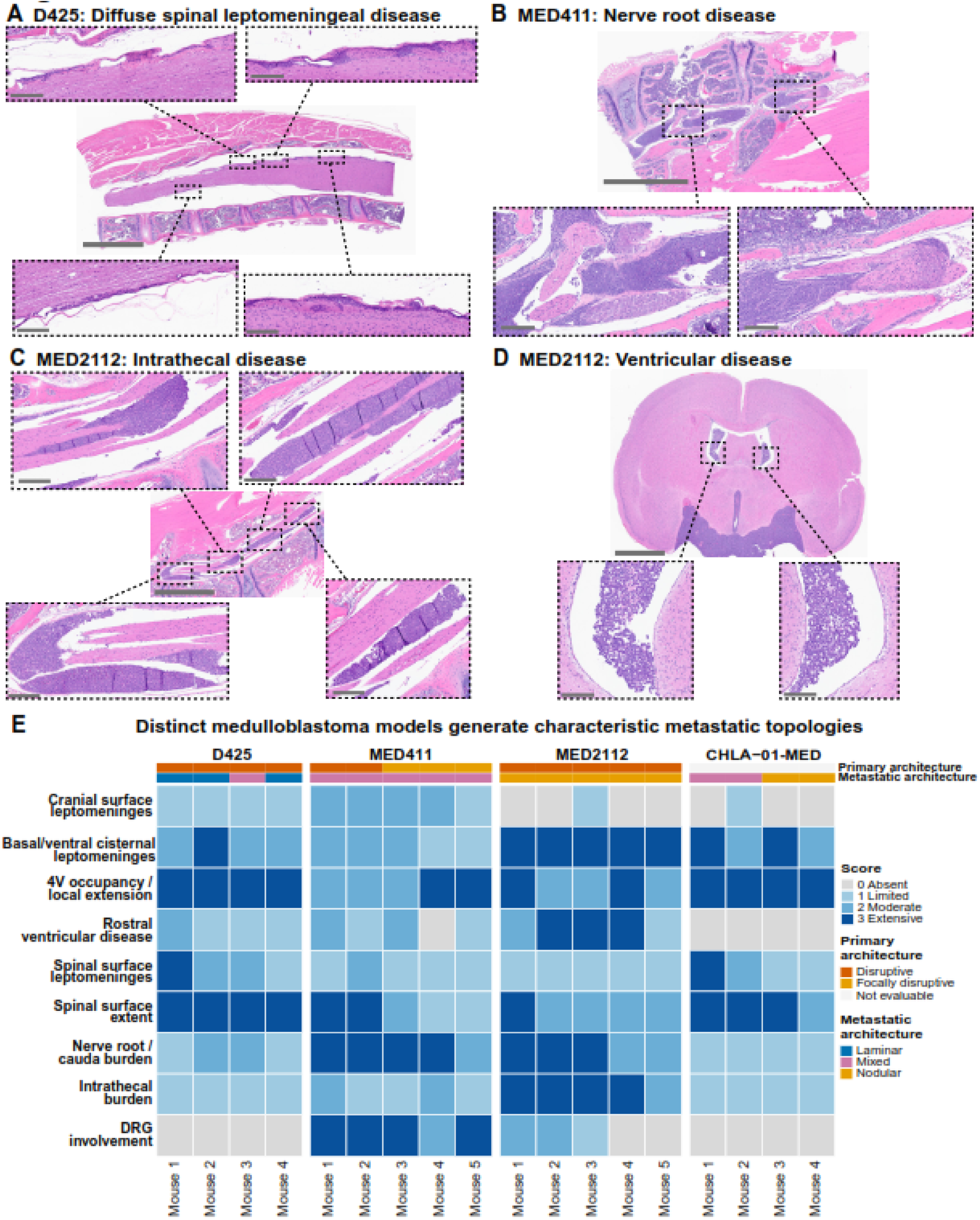
Cross-model comparison of metastasis topology reveals role of tumor biology in dissemination. A-D) Representative histological images of topologies enriched between models. Scale bar = 2 mm. Inset scale bar = 200 µm. E) Heat-map illustrating metastatic topology scores by model. Of note, CHLA-01-MED architecture scored as not assessed (NA) due to lack of parenchymal component of tumors.

To quantitatively assess these differences, we performed multivariate analysis of nine histopathological topology features encompassing cranial, ventricular, and spinal disease. Among mice with confirmed distant metastases (n = 18), metastatic topology differed significantly across tumor models despite standardized midline implantation (PERMANOVA, pseudo-F = 13.57, R² = 0.744, p = 0.0001; Figure 5E). These differences persisted after excluding fourth-ventricular tumor burden, which may reflect contiguous primary tumor extension rather than distant dissemination (pseudo-F = 16.83, R² = 0.783, p = 0.0001), without detectable differences in multivariate dispersion (PERMDISP, p = 0.342). Individual compartment analyses identified significant differences in nerve-root/cauda burden, intrathecal burden, and dorsal-root-ganglion involvement (Holm-adjusted p = 0.0034, 0.0034, and 0.0022, respectively), whereas spinal surface extent did not differ significantly between models (Supplementary Figure 3E–I).

Together, these findings demonstrate that medulloblastoma models arising from a shared anatomical starting location exhibit distinct patterns of craniospinal disease. Thus, primary tumor implantation geometry alone is insufficient to explain metastatic topology, supporting a contribution from tumor-associated biological characteristics to the anatomical distribution of disseminated disease.

## Discussion

Despite substantial advances in understanding the molecular drivers, developmental origins, and clinicopathological characteristics of medulloblastoma, metastatic progression remains a major contributor to treatment failure and mortality. Consequently, contemporary treatment must address the possibility of occult disease throughout the craniospinal axis. Although clinical staging captures the broad extent of dissemination, it provides limited insight into the processes that permit tumor cells to leave the primary site or the factors that govern their subsequent distribution across distinct craniospinal compartments. Addressing these questions requires examining not only the presence of metastatic disease, but also the relationship between primary tumor growth, anatomical boundaries, and the spatial organization of disseminated lesions. In this study, we used orthotopic models and matched craniospinal histopathology to investigate how primary tumor geometry, growth architecture, and tumor-model identity relate to metastatic dissemination and topology. Our findings suggest that metastatic progression is not simply proportional to primary tumor burden or determined by anatomical starting position. Instead, CSF-boundary disruption is strongly associated with dissemination, while distinct medulloblastoma models develop different patterns of craniospinal colonization despite shared implantation geometry.

These findings imply that the development of leptomeningeal dissemination may not be an inevitable consequence of increasing primary tumor burden. Within our experimental models, extensive primary tumors frequently remained confined to their anatomical site, whereas tumors with relatively limited primary burden could develop widespread craniospinal disease. Although these observations cannot completely rule out microscopic dissemination (M0) or define the temporal sequence of metastatic progression, they highlight the potential importance of primary tumor growth characteristics in influencing metastatic behavior. Clinically, patients without detectable metastatic disease represent a heterogeneous population in whom the absence of radiographic or cytologic evidence of dissemination does not necessarily indicate an absence of occult disease or metastatic potential[29]. Understanding the biological features that distinguish localized from disseminating tumors may ultimately improve our ability to identify patients at risk for metastatic progression and inform more individualized approaches to treatment.

An important limitation of this study is that we have not identified the molecular mechanisms responsible for the distinct metastatic topologies observed across medulloblastoma models. Furthermore, because metastatic disease was assessed at neurological endpoint, we cannot determine whether these differences reflect preferential seeding of anatomical compartments, differential survival and expansion following dissemination, or differences in disease duration and progression. Species-specific differences in CSF anatomy and circulation may also influence the distribution of metastatic disease[30, 31]. Notably, unlike humans, mice lack a patent median aperture of the fourth ventricle (foramen of Magendie), with CSF exiting predominantly through the lateral apertures. These anatomical differences may influence the routes by which disseminated tumor cells encounter craniospinal compartments in mice, potentially limiting direct extrapolation of the observed metastatic distributions to patients. Nevertheless, distinguishing the contributions of anatomical access, tumor-cell dissemination, and site-specific colonization requires experimental models that capture the heterogeneity of metastatic disease. By systematically manipulating primary tumor geometry, characterizing the relationship between primary growth architecture and CSF-facing boundaries, and quantifying disease across defined craniospinal compartments, our study establishes an experimental and analytical framework for investigating these processes. The distinct metastatic patterns observed across models provide phenotypic differences that can now be interrogated through targeted perturbation of tumor-cell programs and their interactions with the metastatic microenvironment. Future studies integrating longitudinal assessment of dissemination with comparative analyses of human metastatic disease will be important for determining the mechanisms and clinical relevance of these patterns.

## Funding

This work was supported by grants from CancerFree KIDS (Wolf Pack Research Grant, JBR), St. Baldrick’s Foundation Scholar Award (Miracles in Memory of Michael St. Baldrick’s Scholar, JBR), Alex’s Lemonade Stand Foundation Young Investigator Award (JBR), Hyundai Hope On Wheels Bridge2K (JBR), and Nationwide Children’s Hospital (JBR). LJG is supported by Pelotonia on a Pelotonia Graduate Scholar fellowship.

## Conflict of Interest

The authors declare no financial or personal conflicts of interest related to this work.

## Authorship

L.N.R, L.J.G, E.A.T, D.V.K, R.J.W.R, and J.B.R conceived the study. L.N.R, L.J.G, A.C.G, R.G, A.M.A, K.P performed experiments. L.N.R and L.J.G wrote initial manuscript. L.N.R, L.J.G, R.J.W.R, and J.B.R edited manuscript. R.D.R provided reagents. All authors read and approved the final version of the manuscript.

## Data Availability

The data that support this study are available at reasonable request to corresponding author (J.B.R).

## Acknowledgements

We are grateful to members of the laboratory of Dr. Ryan Roberts for valuable discussions and critical reading of this manuscript. We also acknowledge the Histopathology and Microscopy Core facilities at Nationwide Children’s Hospital Abigail Wexner Research Institute for their excellent technical expertise. The authors acknowledge the use of ChatGPT5, solely for text editing and clarity. All schematics in this manuscript were generated using BioRender.

## Notes

### Competing Interest Statement

The authors have declared no competing interest.

