## Supplemental Figures for "CSF Boundary Disruption and Tumor-Dependent Patterns of Medulloblastoma Metastasis"

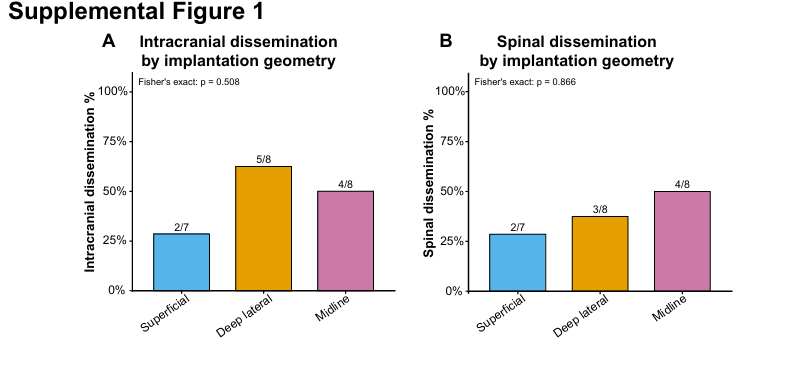
**Supplemental Figure 1:** Effect of primary tumor geometry on dissemination. A-B) Primary tumor geometry does not significantly alter intracranial or spinal dissemination.

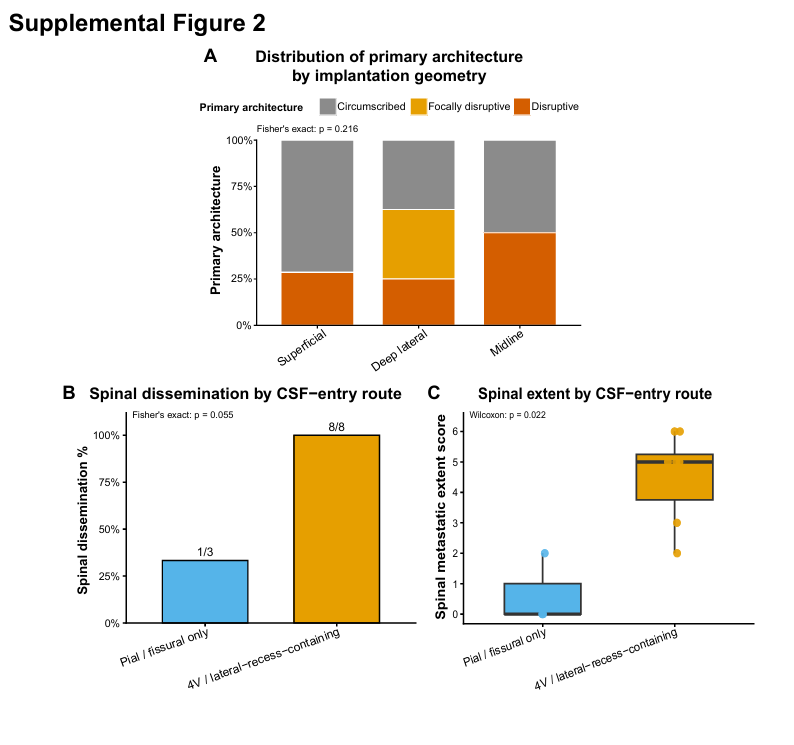

**Supplemental Figure 2:** Effect of tumor geometry on architecture. A) No significant difference in tumor architecture by geometry. B-C) The ventricular CSF breach significantly increased both the frequency and extent of spinal dissemination compared to tumors that disrupted cerebellar pia layer.

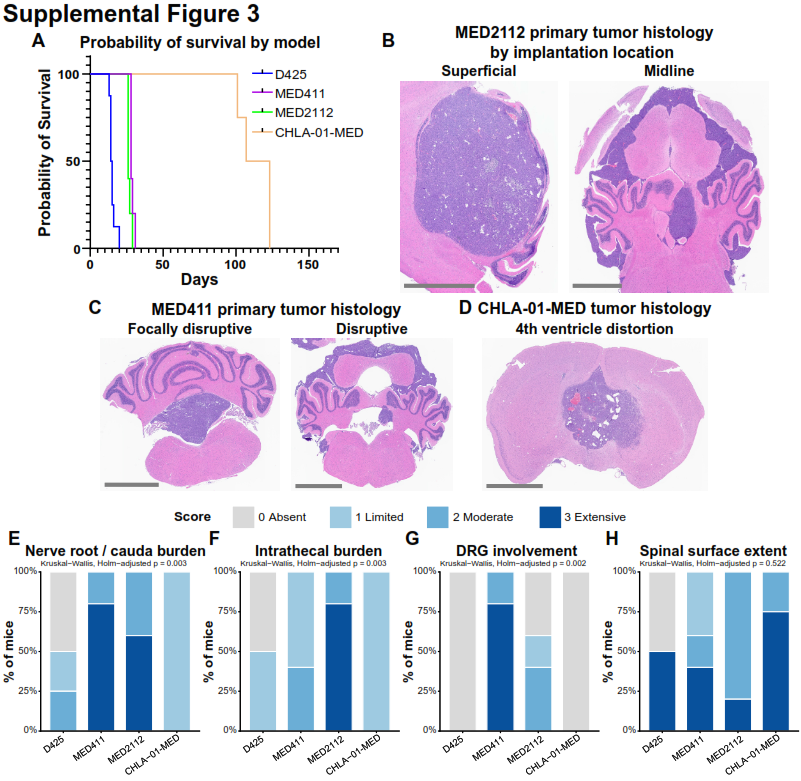

**Supplemental Figure 3:** Model survival and topology A) Kaplan Meier curve shows survival by model. B-D) Representative histological images of primary tumors from MED411, MED2112, and CHLA-01-MED. Scale bar = 2 mm. E-H) Frequency of individual metastatic locations by model.
